# The Role of *SIAMESE* in G2 Checkpoint Regulation in *Arabidopsis thaliana*

**DOI:** 10.1101/2025.05.16.654532

**Authors:** Martha Adele Schwall, Frances Clark, Renee Dale, Adrienne Roeder, Nao Kato, John C. Larkin

## Abstract

In Arabidopsis, SIAMESE (SIM) is a cyclin-dependent kinase inhibitor that restricts progression through mitosis. SIM is well known as a regulator of endoreplication in trichomes and roots. Mathematical modeling of the cell cycle has indicated that SIM may also modulate the length of G2 during mitotic cycles, potentially replacing the WEE1/CDC25 circuit that regulates G2 timing in animals and fungi, which is absent in plants. The predictions of this model were tested in several ways. First, the root meristem is longer in *sim* mutant roots than in wild-type (WT) roots. Second, two independent methods of measuring cell cycle phases, long-term live-cell imaging using fluorescent protein cell cycle markers and 5-ethynyl-2-deoxyuridine (EdU) pulse-chase imaging, showed that the length of G2 in root meristem cortex cells is shorter in *sim* mutant plants compared to WT, consistent with the view that these changes in G2 length are due to greater G2 CDK activity in *sim* mutants. Additionally, the fluorescence of a CYCB:GFP fusion both rose and declined more sharply in the mutant than in wild-type. Because both transcription and degradation of this fusion are directly affected by G2 CDK activity, this result is consistent with *sim* mutants having greater G2 CDK activity. Taken together, these results suggest that, in addition to its known role in inducing endoreplication, SIM may play a role in regulating the length of G2 during mitotic cycles, at least partially replacing the absence of WEE1/CDC25 regulation of the mitotic cell cycle.

## Introduction

The proliferation of individual cells is the basis for all eukaryotic life. Cell division and differentiation are necessary for the formation of tissues and, ultimately, organisms. The cell cycle is highly controlled and most components responsible for control are highly conserved across the eukaryotes. The checkpoints of G1/S and G2/M transitions rely on the activity of Cyclin Dependent Kinases (CDKs), which form heterodimeric complexes with Cyclins (CYCs) with some substrate and checkpoint specificity (Jackman et al., 1995; Pavletich, 1999, Nowack et al., 2012). This canonical view of the cell cycle was developed primarily from work in animal and fungal systems.

Animals and fungi both lie within the larger ancient eukaryotic group known as the opisthokonts. Plants diverged from the opisthokonts early in the evolution of eukaryotes, and thus differences in cell cycle regulation between plants and opisthokonts may give insights into the evolution of the eukaryotic cell cycle(Cross et al., 2011). Plants have a unique class of plant-specific CDKs, known as CDKBs, that function primarily to promote mitosis (Boudolf et al., 2001; Vandepoele et al., 2002; Dewitte and Murray, 2003). In addition to regulation by their CYC partners, CDKs are regulated by CDK-activating kinases and CDK inhibitors (CKIs). Our study focuses on the G2/M checkpoint, which allows for both checks of the integrity of the newly synthesized DNA and preparation for mitosis.

In yeast and animals, G2 timing is controlled via CDK inhibition by WEE1 and subsequent activation by CDC25 (Nurse and Thuriaux, 1980; Nurse et al, 1976), both of which are regulated by Polo-like kinases (PLKs)that control entry into mitosis and functioning of the mitotic machinery (Barr et al, 2004). Despite the conserved nature of the cell cycle and G2/M checkpoint control between eukaryotes, plants have distinctive features that differentiate them from other eukaryotes, particularly with respect to control of G2. Arabidopsis *wee1* mutants have no effect on cell cycle progression of Arabidopsis under ordinary conditions, and although *wee1* mutants are hypersensitive to DNA damaging agents, its role in the DNA damage checkpoint does not involve phosphorylation of CDKA;1 (De Schutter et al, 2007; Dissmeyer et al, 2010). In addition, plants lack identifiable CDC25 and PLK homologs (Bleeker et al, 2006; Kurasawa et al, 2020).

In Arabidopsis, a transcriptional wave of G2/M genes is regulated by a multiprotein complex, similar to the animal DREAM complex, reinforcing CDK activity, microtubule reconfiguration, and chromatin remodeling. Members of the plant DREAM complex are activated by plant-specific B-type cyclin/CDK complexes (Sablowski and Gutierrez, 2022; Polyn et al, 2015). In the absence of the G2 regulation by WEE1, CDC25 and PLK seen in other eukaryotes, it remains unclear what, if any, negative regulators restrain CDK activity to regulate timing of the G2 phase in mitotically dividing plant cells.

The SIAMESE-RELATED (SMR) family of CKIs, of which SIAMESE (SIM) is the founding member, are known to cause endoreplication when expressed at high levels (Churchman et al, 2006, Roeder et al. 2010). Endoreplication, repeated synthesis of DNA without mitosis, plays roles in organ growth, cell differentiation and stress responses (De Veylder et al, 2011; Lang and Schnittger 2020). SIM binds to and inhibits the activity of CDKA;1 and CDKB, which prevents the CDK-driven self-sustaining G2/M transcription and blocks mitosis in trichomes and roots (Churchman et al, 2006; Van Leene et al, 2010; Kumar et al., 2015; Wang et al., 2020; Bhosale et al. 2018). Another member of the SIAMESE-RELATED (SMR) family, SMR1/LGO, regulates endoreplication in leaves, sepals and roots (Roeder et al, 2010; Dubois et al, 2023). Although SIM and other SMRs are presently known as regulators of endoreplication, their ability to inhibit mitotic CDK activity makes them obvious candidates for a role in regulating the length of G2 in mitotic cells.

The arrangement of tissues in the Arabidopsis root is highly structured and has a distinct transition zone from mitotically dividing cells to endoreplicating cells. This makes the root an ideal organ in which to study cell cycle regulation. The root tip contains the stem cell niche, quiescent center (QC), and the meristem, where cells divide mitotically (Dolan et al, 1993, Lavrekha et al, 2017). Moving shootward (proximally), the mitotically dividing cells transition to endoreplication cycles in the elongation zone, and finally, cells mature into their respective tissues in the maturation zone (Bhosale et al, 2019). *SIM* is highly expressed in the elongation zones of roots where it, along with SMR1 and other SMRs, is necessary for the cells to endoreplicate and differentiate (Bhosale et al, 2019; Li et al, 2016). However, *SIM* is also expressed at lower levels in meristematic cells (Li et al, 2016), consistent with a possible role in mitotic cycles. Through modeling, live cell imaging, and EdU labeling, here we show that *SIM* is integral to establishing the length of the G2 phase in the root meristem cortex cells, in addition to its previously known role in endoreplication.

## Results

### Root growth rate and meristem length are greater in *sim* mutants

Among members of the *SMR* family, *SIM* transcripts are the most abundant in the root meristem (Li et al. 2016). For this reason, we investigated the effects of *sim* mutants on root meristem function. We first measured total root length over the course of 9 days after sowing (das). Beginning on 6 das, the *sim* roots were significantly longer than WT (p < 0.001), a trend that continued through the remainder of the growth meristem length at 6 das (p < 0.001) (Figure 1B, 1C), as well as significantly more cells in the meristematic zone (Supplemental Figure 1A). The increased meristem length could be a direct consequence of either a delayed onset of endoreplication or a shorter cell cycle length, leading to a longer meristematic zone with more cells.

**Figure 1.**
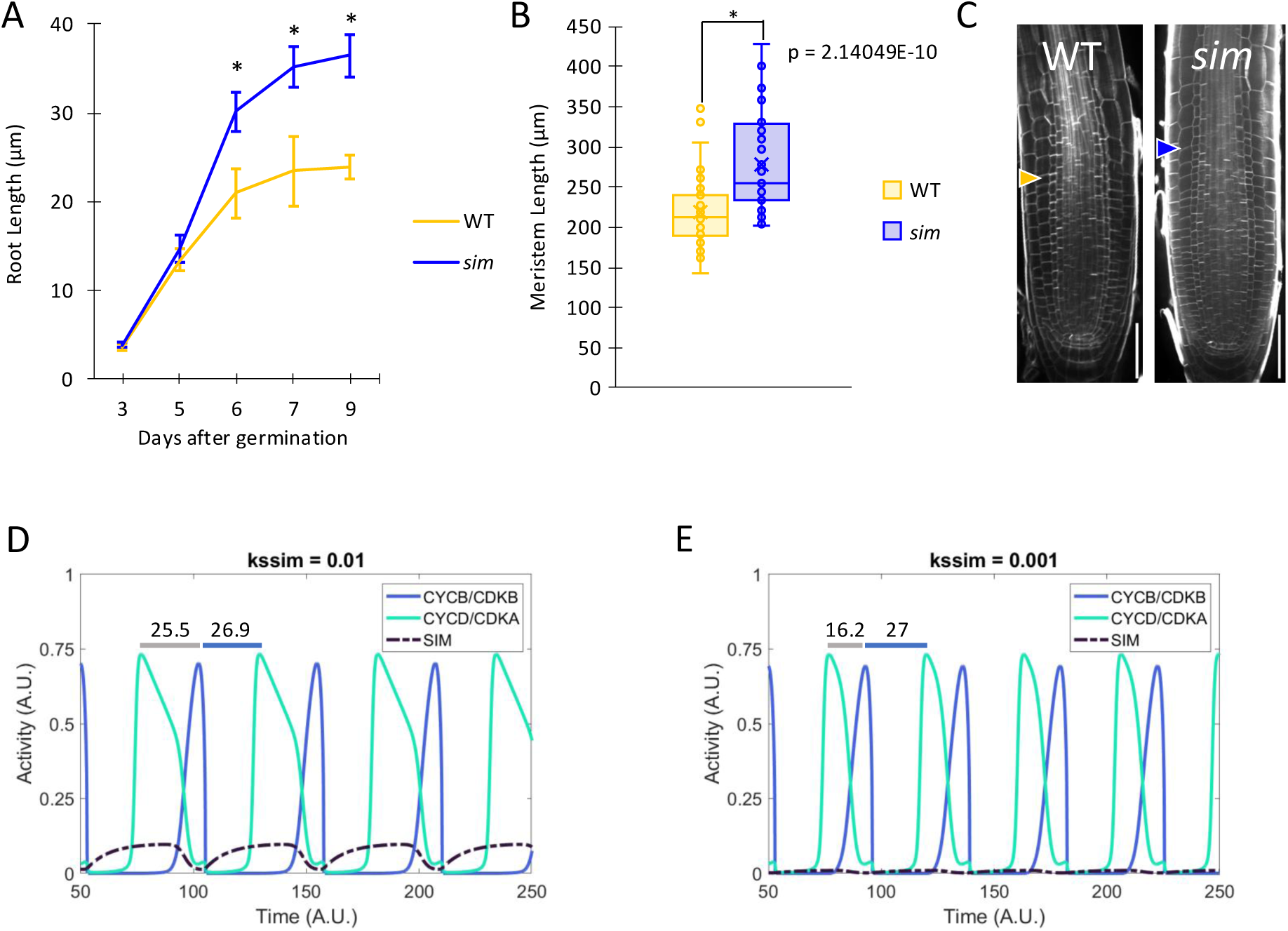
Root growth, meristem size, and model prediction that *sim* mutants have shorter G2 phase and total cell cycle. A. Root growth over time. Days 3 and 5 are not significantly different. *sim* roots are significantly longer on days 6-9 (p = 3.05E-05; p = 6.01E-07; p = 6.69E-13. days 6-9 respectively) (Unpaired students t-Test). WT (n = 106 roots); *sim* (n = 105 roots). B. Meristem length. *sim* has significantly longer meristem (p = 2.14E-10) (Unpaired students t-Test). WT (n = 74 roots); *sim* (n = 48 roots). C. PI-stained images of one representative of WT and *sim*. Arrows indicate meristem boundary. Scale bars = 50 μm. D. Model predictions for CYCB/CDKB and CYCD/CDKA activity at 0.01 kssim. kssim = 0.01 is considered “wild type” *SIM* synthesis. Grey bar indicates duration of S-G2-M. Time in arbitrary units. Blue bar indicates duration of M-G1-S. E. Model predictions for CYCB/CDKB and CYCD/CDKA activity at 0.001 kssim. Grey bar indicates duration of S-G2-M Time in arbitrary units.

### Modeling predicts that SIM synthesis may affect G2 duration

Using a classic cell cycle model modified for the plant cell cycle, Roodbarkelari et al. (2010) found that increasing the rate constant for synthesis of SIM ten-fold could model the transition from mitotic cycles with alternating peaks of G1/S and G2/M CDK activity to endoreplication (CYCD/CDKA and CYCB/CDKB, respectively). We used this model to predict whether SIM also plays a role in the duration of G2 by varying the SIM synthesis rate constant over a range below the threshold needed to drive the model into endoreplication (Figure 1D and 1E, Supplemental Figure 1B-F). Assuming that the peaks of G1/S and G2/M CDK activity mark the start of S-phase and mitotic anaphase, respectively, the estimated lengths of M-G1-S, S-G2-M, and total cell cycle length in arbitrary time units were estimated. Increasing the SIM synthesis rate caused an increase in the S-G2-M time between the G1/S CDK activity peak and the G2/M CDK peak (Figure 1D,E, Supplemental Figure 1B,E). Once the rate of SIM synthesis rate was raised above 0.04, the model predicted entry into endoreplication, i.e. repeated cycles of S-phase CDK activity with no intervening peak of M-phase CDK activity (Supplemental Figure 1C). In contrast, varying the rate of SIM synthesis predicted no effect on the time from the M-phase CDK peak through G1 to the S-phase CDK peak (M-G1-S, Supplemental Figure 1D). The total cell cycle duration (M-M) increased solely by the increased length of the S-G2-M portion of the cycle (Supplemental Figure 1F). Thus, the model predicted a direct correlation between SIM synthesis rate and S-G2-M duration, indicating that the hypothesis that SIM inhibition of G2/M CDK activity plays a role in regulation of the length of G2 during mitotic cell cycles is reasonable.

### Long-term live-cell imaging shows that G2 duration is significantly shorter in *sim* root cortex cells

To test the prediction that reducing the level of SIM function can result in a shorter G2 phase, we introduced the CYTRAP live cell imaging system into WT and *sim* backgrounds. In this system, an RFP fusion protein including the (C3) region of CDT1a expressed from an S phase-specific promoter is produced starting at the beginning of S-phase and is degraded during mid-G2, while a GFP-tagged CYCB1 expressed from its own promoter becomes visible during G2 and is degraded as anaphase begins (Yin et al., 2014). Because previous work indicated that G2 length differs along the distal-proximal axis of the root meristem (Otero et al. 2016), two meristem regions, distal meristem (the 40%of the total meristem length closest to the QC, closest to root tip) and proximal meristem (the 60% of the total meristem length farthest from QC, farther from root tip) were analyzed separately.

Live imaging showed that *sim* distal cortex cells had a significantly shorter S/G2 duration (RFP fluorescence, Figure 2A, p = 0.003) and a shorter G2/M duration (GFP fluorescence, Figure 2B, 2C, p < 0.001) (Figure 2B, 2C Label GFP/RFP phases) compared to WT, and *sim* proximal cortex cells had a significantly shorter G2-M duration (Figure 2B, p = 0.001). When the normalized peak GFP fluorescence intensities of WT and mutant were aligned at their metaphase maxima, it was apparent that both the rise and fall of GFP levels was more rapid in *sim* than in WT (Figure 2D).

**Figure 2.**
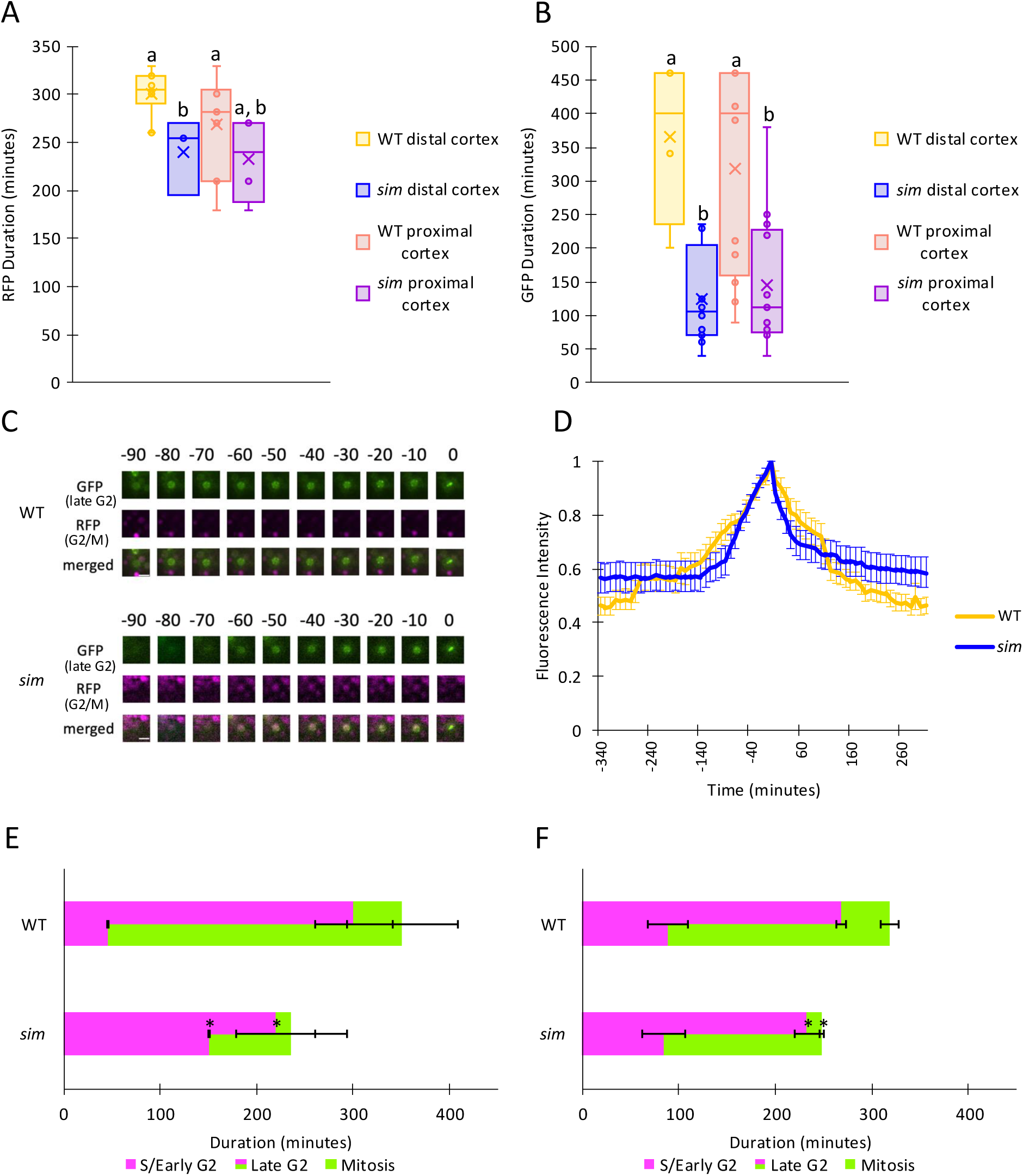
Live imaging using Cytrap markers indicate that *sim* mutants have a shorter G2 than WT. A. RFP fluorescence duration. *sim* distal cortex cells have significantly shorter RFP duration (p = 0.0029) (ANOVA with Bonferroni correction and unpaired students t-Test). WT nuclei (n = 34); *sim* nuclei (n = 25). B. GFP fluorescence duration. sim distal and proximal cortex cells have significantly shorter GFP duration than WT (p = 0.00021 and 0.0012, respectively) (ANOVA with Bonferroni correction and unpaired Students t-test). WT nuclei (n = 16); *sim* nuclei (n = 25). C. Single nuclei tracking of one representative nucleus for WT and *sim*. Time 0 is set as the time when GFP fluorescence is localized to chromosomes on the metaphase plate. D. Fluorescence intensity of GFP expressing nuclei in WT and sim. Time 0 is set as the time of mitosis. Fluorescence intensity normalized to 0 (mean ± s.e.). WT nuclei (n = 16); *sim* nuclei (n = 25). E. Proximal cortex S/Early G2, Late G2, and mitosis durations (mean ± s.e.). WT nuclei (n = 29); *sim* nuclei (n = 17). S/Early G2 (p = 0.441); Late G2 (p = 0.025); Mitosis (p = 0.0013) (ANOVA with Bonferroni correction and unpaired Students t-test). F. Distal cortex S/Early G2, Late G2, and mitosis durations (mean ± s.e.). WT nuclei (n = 21); *sim* nuclei (n = 33). S/Early G2 (p = 3.92e-08); Late G2 (p = 5.46e-19); Mitosis (p = 0.065) (ANOVA with Bonferroni correction and unpaired Students t-test).

When the entire S-G2-M period (RFP and GFP fluorescence combined) is considered, the *sim* mutant progresses through these periods much faster than WT in both the distal and proximal meristem (approximately 115 minutes faster and 70 minutes faster, respectively, Figure 2E, 2F), with most of this reduction being due to a reduced period of GFP expression. These results support the model prediction that *SIM* function affects G2 duration.

### EdU pulse-chase labeling shows G2 duration is significantly shorter in *sim* distal cortex cells

In a second approach that more directly measures differences in G2/M duration, we labeled roots with a short pulse of 5-ethynyl-2’-deoxyuridine (EdU) followed by a chase of varying times in media supplemented with thymidine, as used by others to directly measure G2 duration (Otero et al, 2016; Echevarria et al 2022). The first cells in which EdU fluorescence is detected on the metaphase plate of mitotic cells (mitotic nuclei) will be the cells pulse-labeled at the very end of S phase. The fraction of EdU-labeled mitotic nuclei relative to total mitotic nuclei was quantified, and the time at which 50% of the maximum mitotic nuclei were EdU-labeled was used to estimate G2 duration for each genotype.

By this measure, G2 duration in *sim* distal cortex nuclei was significantly shorter than WT (p = 0.029) by about 94 minutes on average (Figure 3A, 3B). The proximal cortex G2 duration was not significantly different between *sim* and WT, although the mean time for the mutant was 70 minutes less than for WT, a trend suggestive of a shorter G2 (Figure 3A, 3C). Taken together, these results indicate that *SIM* has the greatest impact on G2 in the cells closest to the QC. This reduced time in G2 of the *sim* mutant is similar to the amount of reduction in S-G2-M observed with the Cytrap markers (Figure 2E, 2F).

**Figure 3.**
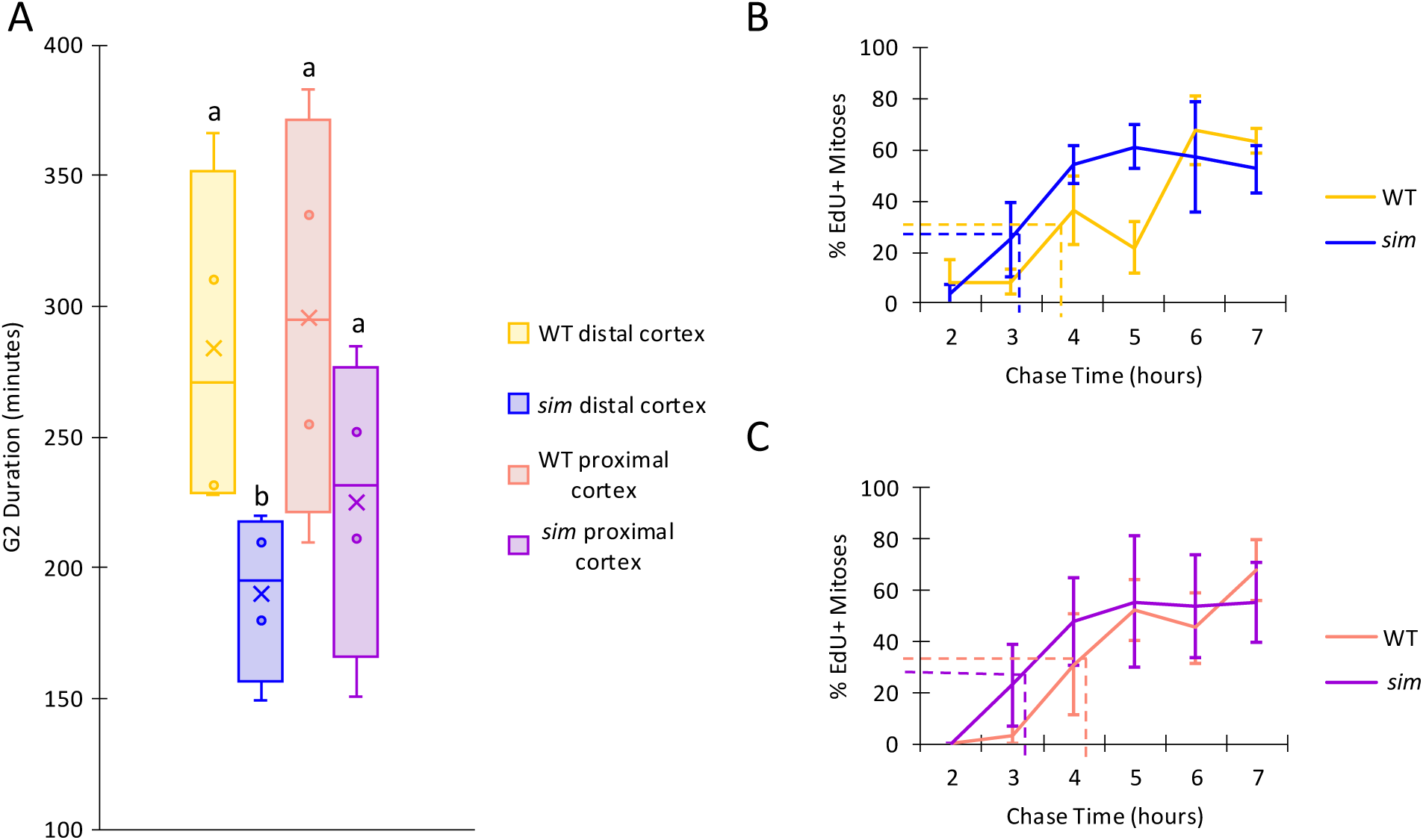
EdU pulse-labeling demonstrates that G2 duration is significantly shorter in sim distal cortex cells. A. G2 duration is significantly shorter in *sim* distal cortex cells (p = 0.0294) (ANOVA with Bonferroni correction and unpaired Students t-test). Data points are G2 duration calculated from 4 replicate experiments. Total number of nuclei analyzed for each genotype were: WT (n= 101 EdU-, n = 121 EdU+), sim (n = 124 EdU-, n = 185 EdU+). Total number of plants analyzed for each genotype were: WT (n = 83); *sim* (n = 84). B. Distal cortex percentage of EdU+ nuclei (mean +- s.e.). Total number of nuclei analyzed for each genotype were: WT (n = 37 EdU-, n = 53 EdU+); *sim* (n = 67 EdU-; n = 113 EdU+).C. Proximal cortex percentage of EdU+ nuclei (mean +- s.e.). Total number of nuclei analyzed for each genotype were: WT (n = 64 EdU-; n = 68 EdU+); *sim* (n = 57 EdU-; n = 72 EdU+).

To confirm these results, we examined a second allele of *sim*, *sim-3*, which has a point mutation changing a proline at position 36 to serine. This proline is conserved in all *SMR* family members (Kumar et al, 2015). EdU pulse-labeling of the *sim-3* allele demonstrated a similar reduction in G2 length of distal cortex cells in the meristem (Supplemental Figure 2A). The *sim-3* allele also has a significantly longer meristem than WT (Supplemental Figure 2B).

### There is no difference in S phase and total cell cycle duration in distal *sim* cortex cells

To estimate the duration of S phase and the total cell cycle, roots were incubated in media supplemented with EdU for a range of times. The percentages of EdU positive nuclei were then measured over time, and the regression line of EdU positive nuclei was used to estimate the durations both the length of S phase and total cell cycle length, assuming linear incorporation of EdU over time (Figure 4A, 4B; Hayashi et al, 2013).

**Figure 4.**
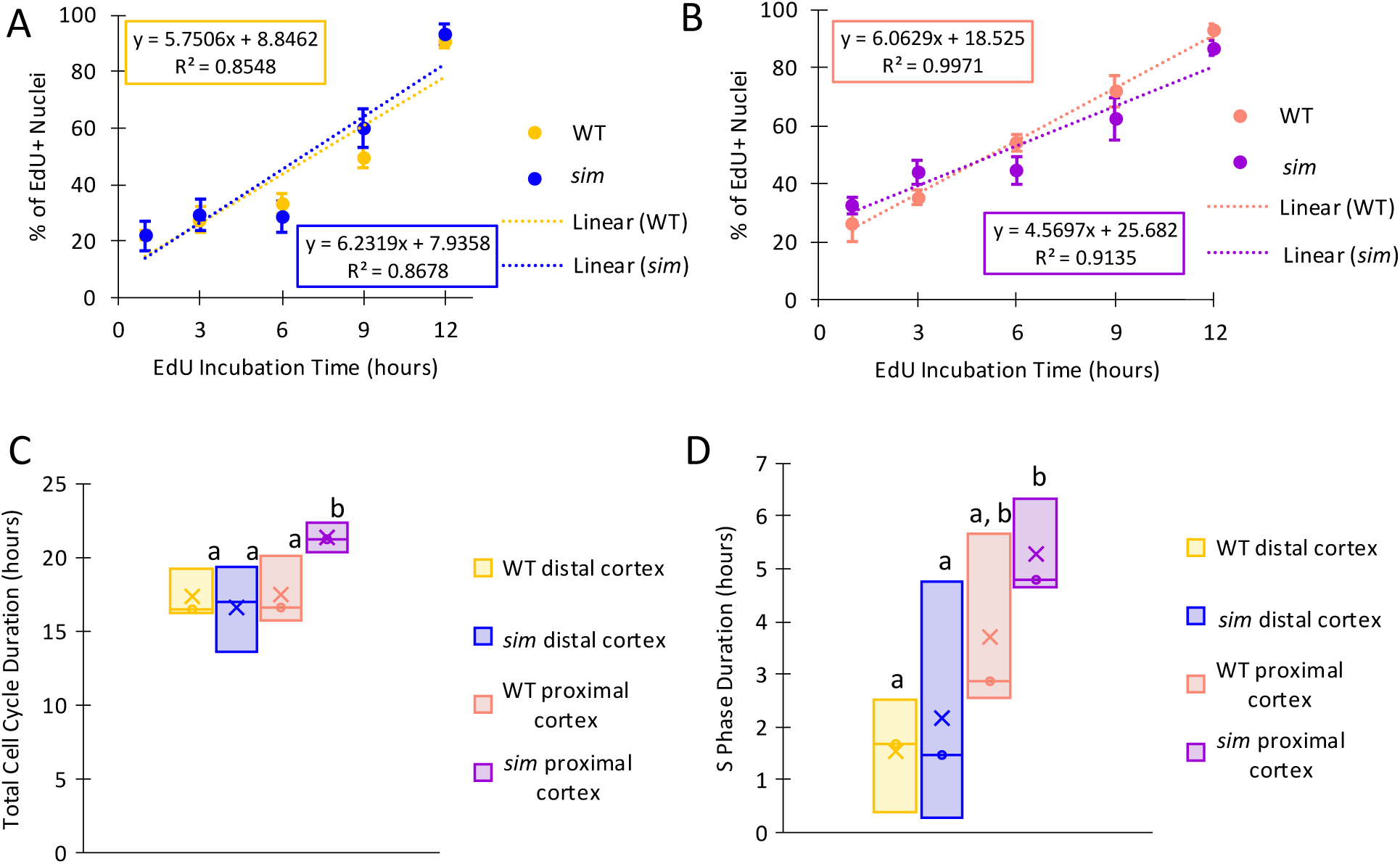
Total cell cycle duration in distal cortex cells is not affected by *SIM*. A. Distal cortex percent of EdU+ nuclei (mean ± s.e.). Linear regression lines are not significantly different (p = 0.465) (Pooled Slopes Test t-Test). Total number of nuclei analyzed for each genotype were: WT (n = 588 EdU-; n = 445 EdU+); *sim* (n = 564 EdU-; n = 451 EdU+). B. Proximal cortex rate of EdU+ nuclei (mean ± s.e.). Linear regression lines are not significantly different (p = 0.063) (Pooled Slopes Test t-Test). Total number of nuclei analyzed for each genotype were: WT (n = 671 EdU-; n = 800 EdU+); *sim* (n = 520 EdU-; n = 577 EdU+). C. Total cell cycle duration. *sim* proximal cortex nuclei have significantly longer total cell cycle than WT and *sim* distal cortex nuclei (p = 0.0429) (ANOVA with Bonferroni correction and Students t-test). Duration was calculated by dividing 100 (maximum EdU+ labeling) by the slope. Figures are mean durations calculated from three replicated experiments. Total number of plants analyzed for each genotype were: WT (n = 51); *sim* (n = 54). D. S phase duration. *sim* proximal cortex nuclei have significantly longer s phase than *sim* distal cortex nuclei (p = 0.0495) (ANOVA with Bonferroni correction and unpaired Students t-test). Duration was calculated by dividing the y intercept by the slope. Points on graph are duration calculated from each replicate. The mean is the average of three replicates. Figures are mean durations calculated from three replicated experiments. Total number of plants analyzed for each genotype were: WT (n = 51); *sim* (n = 54).

In distal cortex cells closer to the QC, there was no significant difference in total duration of the cell cycle between WT and *sim* (Figure 4C), while total cell cycle duration was significantly longer in *sim* proximal cortex nuclei than WT proximal cortex nuclei (p = 0.043),. When comparing cell cycle lengths of distal and proximal cortex cells within each genotype, the total cell cycle length was significantly longer in *sim* proximal cortex cells than in *sim* distal cortex cells (p = 0.028), a trend that was not observed in WT (Figure 4C). S phase duration was not significantly different between the genotypes in either distal or proximal cortex cells (Figure 4D). Thus, comparing these results with the results in Figures 2 and 3, it appears that loss of *SIM* function primarily affects the length of G2/M rather than other cell cycle phases, at least in distal cortex cells.

### Distal cortex cells of *sim* are significantly smaller than WT

The relationship between cell size and G2 duration is straightforward in yeast, where cells that go through G2 more quickly are smaller (Nurse et al, 1976), and the most straightforward expectation is that plant meristem cells will behave similarly. However, this relationship in plant cells is not well understood (Beemster et al, 2002; Yamada et al, 2022). To investigate the relationship between cell cycle and cell size, we measured the cell volume of cortex cells in WT and *sim* meristems (Fig. 5A,B,C). Distal cortex cells of *sim* meristems had significantly less volume than distal cortex cells of WT (777±82 s.e. μm^3^ vs. 1629±259 s.e. μm^3^, p=0.002, Fig. 5C).

**Figure 5.**
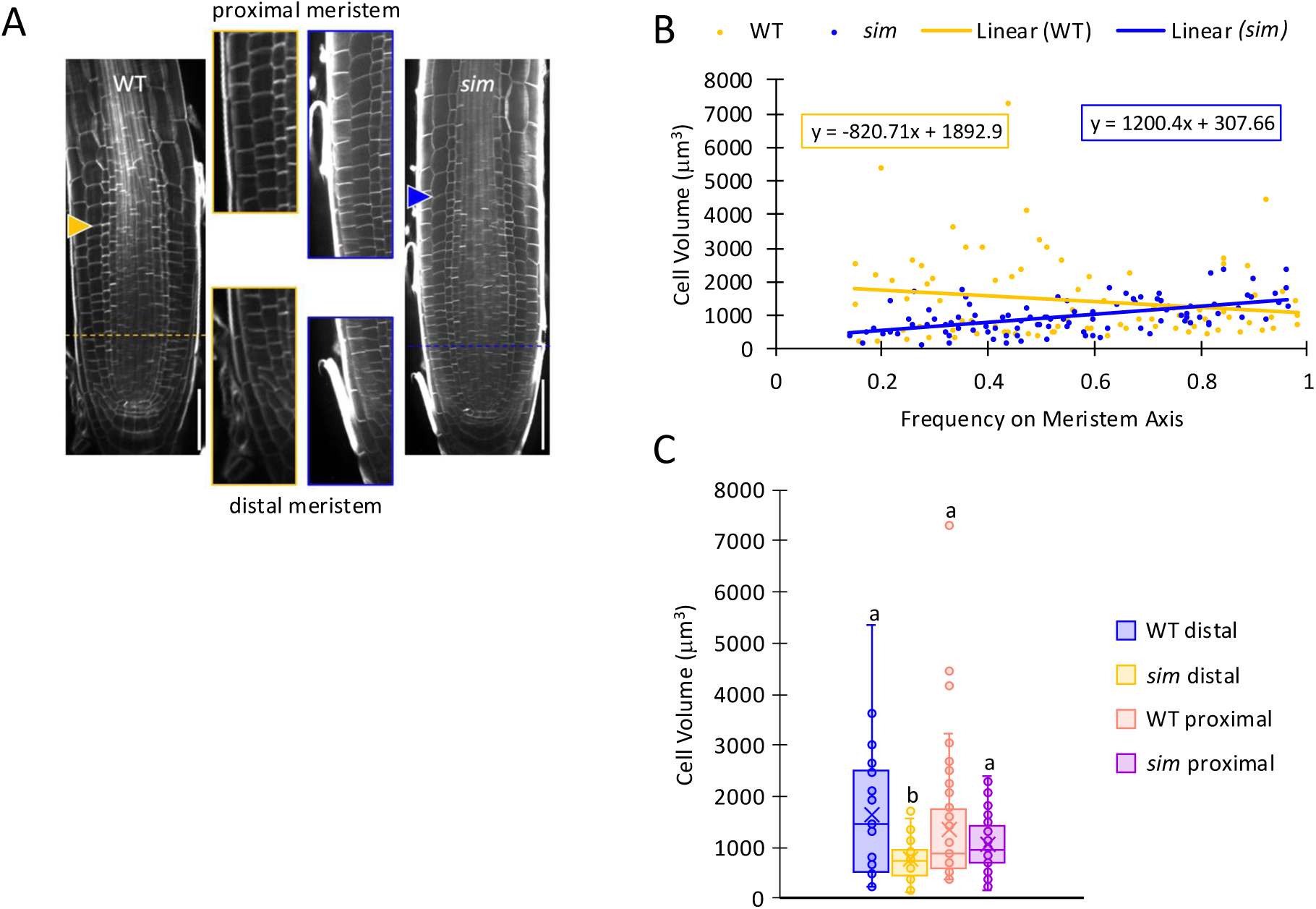
*sim* distal cortex cells are significantly smaller than WT. A. Calcofluor-white stained images of one representative of WT and *sim*. WT (74 roots); *sim* (48 roots). Arrows indicate meristem boundary. Dotted lines indicate distal cortex boundary. B. Cell volume in WT and sim meristem cortex cells mapped to their location along the root meristem. WT (n = 85); *sim* (n = 100). Frequency is defined as the cell distance from the QC divided by the whole meristem length with 1 being the maximum meristem boundary. C. Cell volume in WT and *sim* meristem cortex cells. *sim* distal cortex cells are significantly smaller than WT distal and proximal cortex cells (p = 0.00194; p = 0.000984, respectively) (ANOVA with Bonferroni correction and unpaired Student’s t-test). WT distal cortex (n = 25 cells); *sim* distal cortex (n = 29 cells); WT proximal cortex (n = 60 cells); *sim* proximal cortex (n = 71 cells). Cell walls stained with Calcofluor-white and images analyzed in MorphoGraphX.

## Discussion

### *SIM* plays an important role in the timing of the G2-M interval in the root apical meristem

Mathematical modeling of the cell cycle strongly predicted that SMR-family CDK inhibitors might be able to establish a G2 delay in the cell cycle (Figure 1D, E). As the most highly expressed member of the *SMR* family in the root meristem (Li et al, 2016), *SIM* was an obvious candidate for playing such a role. By two different methods, we have found that *sim* mutant root apical meristem cortex cells progress more quickly from S-phase to mitosis than wild-type root meristems. These two methods are distinct in the way they measure cell cycle phases. In the Cytrap cell cycle marker system, the initiation of RFP expression at the beginning of S phase by transcription from the S phase-specific HTR2 promoter and the termination of GFP-CYCB1 expression at anaphase by the APC/C are well-defined events (Yin et al.2014), but the points at which RFP expression is lost or GFP expression becomes visible are not as clearly tied to specific cell cycle events. However, the rise in GFP level depends on transcription from the CYCB1;1 promoter and fall of GFP expression depends on destruction of the fusion protein by the APC/C, both events that are expected to be correlated with mitotic CDK activity. The observation that GFP fluorescence both rose and fell more quickly in *sim* meristem cells than in WT (Figure 2D) strongly suggested that faster progression through G2 in the mutant was due to a more rapid increase in CDK activity, as expected for a CKI mutant.

In contrast, EdU pulse-labeling gives a direct test of the time between the end of DNA synthesis and mitosis. For the distal meristem, these two different methods gave similar estimates for the degree to which the S-G2-M interval is shortened in *sim* compared to WT (Figure 2E,F, Figure 3A), confirming that most of the observed difference was due to faster progression through G2 to mitosis. In the proximal meristem, the situation was less clear. The Cytrap markers showed a significantly shorter S-G2-M interval in the proximal meristem, while the EdU pulse-labeling showed an almost identical shortening, though it did not reach significance. We also found that the volume of *sim* mutant meristematic cells was less than that of WT, significantly so for cells in the distal meristem (Figure 5A), indicating that mutant cells divide at a smaller size. Taken together, these experiments demonstrate that *SIM* plays a significant role in the timing of the G2-M interval, particularly in the distal meristem, as predicted by mathematical modeling (Figure 1D, E).

### *SIM* has little effect on the length of G1/S or total cell cycle duration

Total cell cycle duration for meristematic cortex cells has been reported to be between 23 hours (Rahni and Birnbaum, 2019) and 17.1 hours (Hayashi et al, 2013). S phase duration has been reported to be 2.9 hours (Hayashi et al, 2013). The WT results reported in this study are similar to previously reported findings (Figure 4C, 4D). S phase and total cell cycle length did not differ significantly between the genotypes in distal cells (Figure 4C, 4D), but in proximal cortex cells the total cell cycle time was slightly but significantly longer than in WT (Figure 4C). This was not predicted by our mathematical model. However overall, the *sim* mutation appears to have minimal effects on total cell cycle duration or S phase duration. Although we would expect the total cell cycle for the distal cortex to be about 1.5 hours shorter in *sim* than in WT, due to the decreased length of G2 (Figure 2E, Figure 3A), this difference is within the range of variation in the individual experiments and may not be detectable (Figure 4A, 4B).

### Diverse roles of *SMRs* in the cell cycle

SIM and other SMRs were initially characterized for their roles in establishing endoreplication (Churchman et al, 2006, Roeder et al, 2010, Kumar et al, 2015) and SIM is known to inhibit both CDKA;1 and CDKB1;1 (Kumar et al, 2015; Wang et al, 2020). Recent studies have also implicated several other SMRs under the direct transcriptional control of the transcription factor SCL28 in modulation of the length of G2 and cell size (Nomoto et al, 2022; Yamada et al, 2022; Goldy et al, 2023). However, *SIM* transcripts are more abundant in the meristem than those of any other *SMR* (Li et al, 2016), and *SIM* is not a transcriptional target of SCL28 (Nomoto et al, 2022). Two *SMR*s, *SMR5* and *SMR7*, are induced as direct targets of the G2 DNA damage transcription factor SOG1, blocking cell cycle progression and inducing endoreplication (Yi et al, 2014; Pedroza-Garcia et al, 2021). In contrast to other SMRs, which appear to exclusively inhibit progression through G2, SMR4 was recently shown to extend G1 during asymmetric divisions of stomatal development (Han et al, 2022).

G2 is an important phase of the cell cycle, allowing cells to prepare for mitosis, check the integrity of the newly replicated DNA, and ensure successful proliferation. In animals and fungi, WEE1, CDC25, and PLKs regulate entry into mitosis, and thus the length of G2 (Nurse, 1976; Barr et al, 2004; de Gooijer et al,2017), but this regulatory circuit is absent in plants. Transcription of mitotic cyclins and CDKs and other mitotic components by the plant DREAM complex are the main positive drivers of the G2/M transition in Arabidopsis (Sablowski and Gutierrez, 2022), but the identity of negative regulators of G2/M progression during mitosis, as opposed to endoreplication has remained unclear. SIM and other SMRs are regulated transcriptionally in response to various abiotic and biotic signals (Yi et al, 2014) and are likely also subject to postranscriptional regulation (Dubois et al, 2018; Kumar et al, 2018). Based on the work presented here and the work of others, SIM and other SMRs are well-positioned to flexibly regulate G2 progression either via subtle modulation of the speed of progression to mitosis or by blocking division altogether and promoting endoreplicaton. Additionally, understanding the role of SMR-type CDK inhibitors in the timing of G2 in plants contributes to our understanding of the diversity of cell cycle regulation across the eukaryotes.

## Materials and Methods

### Plant materials and growth conditions

Plants were grown on soil as previously described (Larkin et al, 1999). The Col-0 ecotype of Arabidopsis thaliana was used as the WT control for all experiments. The *sim-1* (referred to in text as “*sim*”) and *sim-3* alleles have been previously described (Churchman et al, 2006). Both alleles were isolated in a Col-0 background and have been backcrossed to Col-0 three times before selfing and recovering homozygotes for experimental use. The *sim-1* allele is point mutation changing the putative At5g04470 start codon from ATG to ATA. The *sim-3*: contains C **→** T mutation, resulting in a Pro-to-Ser amino acid change at position 36. This proline residue is conserved in all *SMR*s (Kumar et al., 2015). After visual identification of homozygotes via their multicellular trichome phenotype, the *sim-1* and *sim-3* allele genotypes were confirmed via PCR using primers as described in Kumar et al, 2015. The Cytrap fluorescent cell cycle marker line was a kind gift from the Umeda Lab (Nara Institute of Science and Technology, Japan). This line was crossed to *sim-1* mutants, the F1 were selfed, and F2 plants were genotyped by visual identification of the *sim* trichome phenotype, followed by PCR genotyping and confirmation of expression of the two Cytrap fluorescent markers.

For root microscopy and growth analysis, seeds were grown on ½ MS plates as follows: Seeds were sterilized in sterilizing solution (50% bleach, 50% water, Tween-20) for 5 minutes, sterilized in 70% ethanol for 1 minute, then rinsed four times with deionized water. The seeds were sown on half-strength Murashige and Skoog medium (Murashige and Skoog, 1962) supplemented with 1% sucrose and 0.8% plant tissue culture agar (Sigma). The plates were placed vertically in a growth chamber kept at 22°C in continuous light. For root growth analysis, beginning three days after germination, roots were imaged every day and primary root growth was measured in FIJI 2.9.0.

### Cell cycle model

The plant-specific cell cycle model used was from Roodbarkelari et al, 2010. Model simulations were run in Matlab 2021a. Code, including parameter values, is given in Supplemental Materials.

### Meristem and cell size analysis

Cell walls were stained according to protocols based on Lu et al, 2021 and Kurihaha et al, 2015. Seedlings 6 days after germination were fixed with 4% paraformaldehyde for 1 h at 23°C. Then the seedlings were washed twice with 1 x PBS and incubated in Clearsee solution (10% xylitol, 15% sodium deoxycholate, and 25% urea in water, Kurihara et al, 2015) for 4 days. Seedlings were then stained with 0.01% Calcofluor White for 1 hour and washed once in Clearsee. Seedlings were mounted on slides in Clearsee.

Roots were imaged on Olympus SpinSR10 Spinning Disk confocal equipped with the Confocal Scanner Unit CSU-W1 SoRa (Yokogawa) at 40x magnification. 0.1 μm Z-stacks of the whole root (about 90 um) were taken of each root. 405 nm laser was used at 80%-100% intensity for visualization of fluorescence.

Images were analyzed with FIJI 2.9.0 (Schindelin et al., 2012). For analysis of meristem length, the meristem was defined as the distance from the QC to the first elongated cell (Baskin et al, 1995). For analysis of cell volume, images were analyzed with MorphoGraphX 2.0.1 (Barbier de Reuille et al, 2015). CNN was generated and ITK watershed autofill was used to create a mesh of the cells (Eschweiler et al, 2019). Volume was calculated from the mesh.

### Long-term live-cell imaging

Roots 6 days after germination were used for imaging. Roots were placed in a chamber made of poly(dimethylsiloxane) gum, and ½ MS +1% sucrose agar slab, and perflurodecalin during imaging (Kirchelle and Moore, 2017). Cytrap roots were observed for 8-10 hours via Olympus SpinSR10 Spinning Disk confocal equipped with the Confocal Scanner Unit CSU-W1 SoRa (Yokogawa) at 20x magnification. 1 μm Z-stacks of the top half of the root (about 50 μm) were taken every ten minutes to minimize photobleaching and unintended root death. 488 nm laser at 10%-30% intensity and 561 nm laser at 5%-20% were used to visualize fluorescence.

For analysis, fluorescence intensity was measured for cells that had trackable fluorescence through the time-lapsed images. Tissue and distance from QC were also recorded for each tracked cell. Images were analyzed with the TrackMate plugin in FIJI 2.9.0 (Ershov et al 2022; Tinevez et al, 2017) and MorphoGraphX 2.0.1 (Barbier de Reuille et al, 2015). For figure 2D, mean fluorescence of all cells was used and normalized to 1.

### EdU pulse chase

Seedlings 6 days after germination were placed in liquid medium (½ MS, 1% sucrose, 10 μM EdU in the Click-iT component A (Invitrogen)) and incubated with EdU at 20^0^C under continuous light for 15 minutes (pulse period). Seedlings were then placed in liquid media ((½ MS, 1% sucrose, 50μM thymidine) for 2-7 hours. EdU was detected following the manufacturers’ instructions for Click-iT with 647 (far-red) azide and stained with Hoechst 33342 for 30 minutes. After 2 washes in PSB, samples were mounted in PBS on slides. Samples were observed via Olympus SpinSR10 Spinning Disk confocal equipped with the Confocal Scanner Unit SCU-W1 SoRa (Yokogawa) at 40x magnification. 0.5 μm Z-stacks of the entire root (about 90 um) were taken of each root. 647 and 405 nm lasers at 80%-100% intensity were used to visualize fluorescence.

For analysis, every mitotic cell (defined by the arrangement of the chromosomes on the metaphase plate) was counted, and its tissue, EdU labeling status, and distance from the QC was recorded. Images were analyzed in FIJI 2.9.0 and MorphoGraphX 2.0.1 (Barbier de Reuille et al, 2015).

Five roots of each genotype (WT and sim) were anayzed per timepoint in each of 4 experimental replicates.

### Long term EdU incubation

Seedlings 6 days after germination were placed in liquid medium (½ MS, 1% sucrose, 10 μM EdU in the Click-iT component A (Invitrogen)) and incubated with EdU at 20°C under continuous light for 1 to 12 hours. EdU was detected following the manufacturers’ instructions for Click-iT 647-azide and stained with Hoechst 33342 for 30 minutes. After 2 washes in PBS, samples were mounted in PBS on slides. Samples were observed on Olympus SpinSR10 Spinning Disk confocal equipped with the Confocal Scanner Unit SCU-W1 SoRa (Yokogawa) at 40x magnification. 0.5 μm Z-stacks of the entire root (about 90 μm) were taken of each root. 647 and 405 nm lasers at 80%-100% intensity were used to visualize fluorescence.

For analysis, all nuclei were counted and tissue and distance from QC to each nucleus was recorded. The ratio of EdU- to EdU+ nuclei was also recorded for each individual root within the tissue types. Images were analyzed in FIJI 2.9.0 and MorphoGraphX 2.0.1 (Barbier de Reuille et al, 2015). 5 roots of each genotype (WT and *sim*) were done per timepoint in 3 experimental replicates.

S and G2 duration were calculated as in Hayashi et al (2013). To calculate S phase and G2 duration, the average ratio of incorporated EdU (EdU-incorporated nuclei divided by Hoechst 33342-stained nuclei, then multiplied by 100) for each timepoint was plotted against the EdU incubation time for cells in the meristematic zone. The averages of each experimental replicate were then averages, and using linear regression, the time for all nuclei to incorporate EdU was estimated by the linear approximation formula (Hayashi et al, 2013):

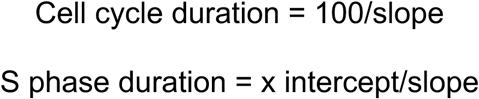

### Statistical methods

To determine statistical significance, ANOVA (XLSTAT plugin for Excel (Excel v 16.89 (24090815)) with Bonferroni correction, SlopesTest and student’s t-Test (Excel v 16.89 (24090815)) were used.

## Accession numbers

*SIAMESE*, AT5G04470.

## Acknowledgements

This work was supported by the National Science Foundation awards NSF-MCB-161572, NSF-BSF-IOS-EDGE 1923589/2019610, NSF-IOS-2014300. We also acknowledge the expert technical assistance of the LSU Shared Instrument Facility, the kind gift of the Cytrap Arabidopsis seed stocks by Dr. Masaaki Umeda of the Nara Institute of Science and Technology, Japan, and critical reading of the manuscript by James Moroney.

## Author Contributions

M.A.S designed experiments, performed experiments and analyzed data, F.C. performed experiments and contributed to writing, R.D. performed computational simulations, A.H.K.R. designed experiments and wrote manuscript, N.K. designed experiments and performed computational simulations, and J.C.L. designed experiments and wrote the manuscript.

## Supplemental Figures

**Supplemental Figure 1.**
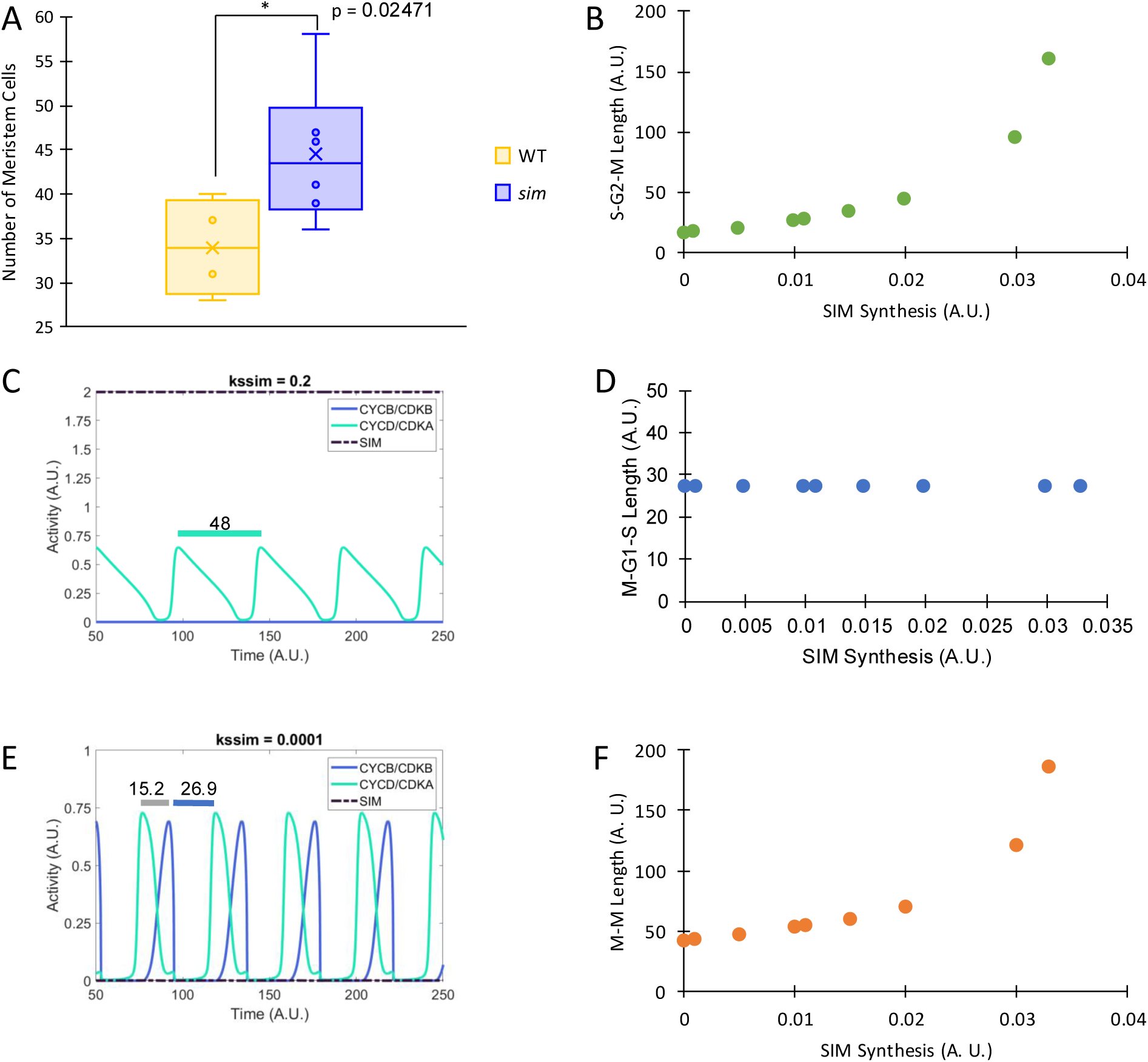
Number of meristem cells and additional model predictions. A: Meristem size by number of cortex cells in meristem (p = 0.0247) WT (n = 117); *sim* (n = 74) (unpaired Students t-test). B: S-G2-M duration dependent on SIM synthesis. Time in arbitrary units. C: Model predictions the cell will go through endoreplication cycles at 0.2 kssim. Time in arbitrary units. D: M-G1-S duration dependent on SIM synthesis. Time in arbitrary units. E: Model predicts the cell will go through mitosis at kssim = 0.0001. Time in arbitrary units. F: Total cell cycle duration dependent on SIM synthesis. Grey bar indicates S-G2-M duration. Time in arbitrary units.

**Supplemental Figure 2.**
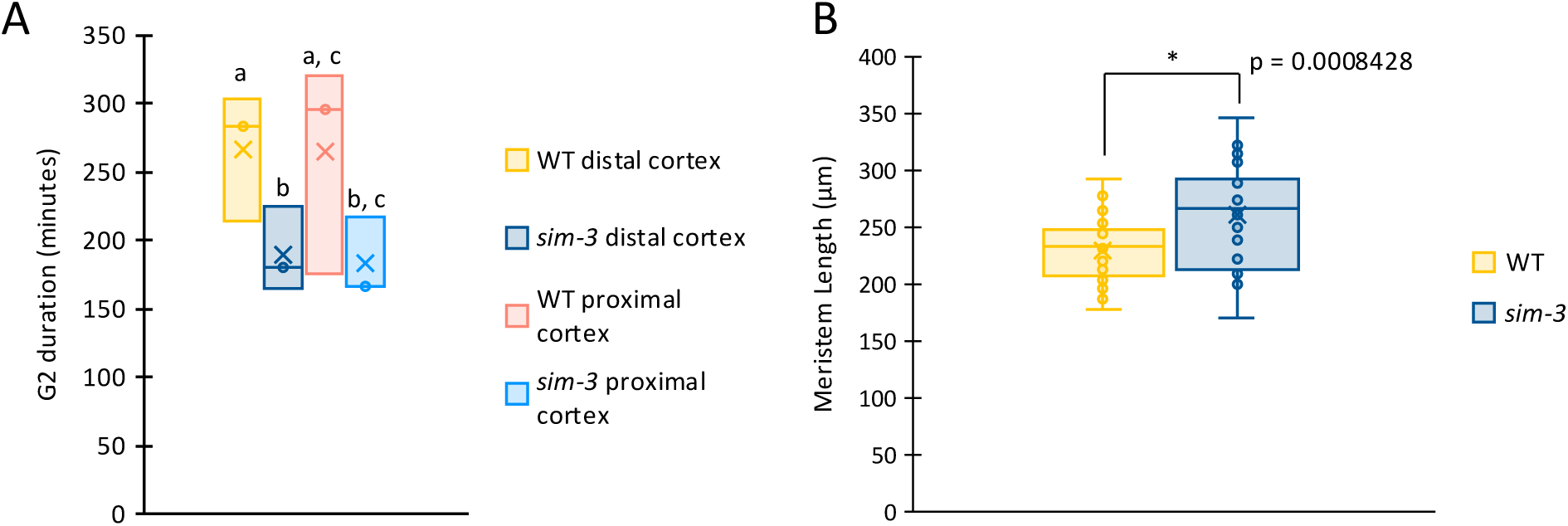
*sim-3* roots have significantly longer meristems and shorter G2 in distal meristem cortex cells than WT. A. Meristem length. *sim-3* has significantly longer meristem (p =0.000842) (unpaired Students t-test). WT (n = 35 roots); *sim-3* (n = 24 roots). B. G2 duration is significantly shorter in *sim-3* distal cortex cells (p = 0.0384) (ANOVA with Bonferroni correction and Students t-test). Data points are G2 duration calculated from 3 replicates of experiment. Total number of nuclei analyzed for each genotype were: WT (n= 93 EdU-, n = 133 EdU+), *sim*-3 (n = 123 EdU-, n = 160 EdU+). Total number of plants analyzed for each genotype were: WT (n = 30); *sim-3* (n = 31). C. Distal cortex percentage of EdU+ nuclei (mean ± s.e.). Total number of nuclei analyzed for each genotype were: WT (n = 40 EdU-, n = 67 EdU+); sim-3 (n = 57 EdU-; n = 64 EdU+). C. Proximal cortex percentage of EdU+ nuclei (mean ± s.e.). Total number of nuclei analyzed for each genotype were: WT (n = 53 EdU-; n = 66 EdU+); *sim-3* (n = 66 EdU-; n = 96 EdU+).

## Supplemental Materials

### Matlab Code to Run the Cell Cycle Model from Roodbarkelari et al. (2010)

The code below is an implementation of the code for the model of Roodbarkelari et al. (2010), formatted to run in Matlab 2021a. Minor changes that do not affect the behavior of the model itself include renaming of model objects to bring them closer to the commonly used names of proteins functioning in the plant cell cycle (the original model was derived from a yeast model and retained yeast naming conventions in many places).

<u>Model code:</u>

function Roodbarkelari_PNAS_2010_code

% code used to run Roodbarkelari et al, PNAS 2010 model

% original model DOI: https://doi.org/10.1073/pnas.1006941107

current=’simhi’;

figure(1)

clf

%% SIM

kssim=.1; % low: 0.01 hi: 0.1 sim-: 0.0001

%% CCS52

ccs52ko=1; % 1 for normal; 0 for knockout

cdhs=1; % cdh synth, 1 for normal, 2 for oe

%% CYCB

k11=0.01;%% cycb: 0.01 normal; 0.03 OE

%% KRP

ko=.2;% k11 -- KRP synth .2 normal, 0.1 KO, 0.3 OE, 0.4 OE++

%% CYCD3

k1311=0.05;% cycd3 mutants: 0.05 normal; 0.075 OE; 0.03 ko

cycd3ko=1; % 1 for normal; 0.5 for knockut

%% other parameters

% original Roodbarkelari parameter names commented; in general, “i” ending

% changed to “1”

k11=0.01;%%k1i

k1=0.1;

lp=1000.0;

lm=1.0;

k31=0.0;

k311=100.0;

j3=0.01;

k41=.3;

k4=40.0;

j4=0.01;

k7=1.0;

j7=0.01;

k8=0.2;

j8=0.01;

k9=0.1;

j9=0.01;

k10=0.04;

j10=0.01;

ko211=0.0;%k21i

ko21=10.0;%k21

jtfb=1.0;

k22=0.5;

k151=0.25;

j15=0.1;

k161=0.01;

k1611=2.0;

j16=0.1;

k131=0;

k141=0.02;

k14=1.0;

lcm=1.0;

lcp=400.0;

k21=0.05; %k2i

k211=100.0;

k2111=1.0;

k12=0.2;

k121=2.0;

k1211=10.0;

kdsim1=0.1;

kdsim=1.0;

% initial conditions as in Roodbarkelari et al PNAS 2010

Xo=[.6 .578 .225 0 .06 0 .03 .923 .275 .085 .06];

tspan=[0:.1:1000];options=odeset(’RelTol’,1e-6);

[t,x]=ode15s(@larkinpnas,tspan,Xo,options,k11,k1,lp,lm,kssim,k31,k311,j3,k41,k4,j4,k7,j7,k8,j8,k9,j9,k10,j10,ko,ko211,ko21,jtfb,k22,k151,j15,k161,k1611,j16,k131,k1311,k141,k14,lcm,lcp,k21,k211,k2111,k12,k121,k1211,kdsim1,kdsim,cdhs,ccs52ko,cycd3ko);

figure(1)

clf

colormap(’jet’)

% skip the burn-in to reduce influence of specific initial condition

% choices

plot(t(5000:10000),x(5000:10000,2),’LineWidth’,2); hold on % MPF

plot(t(5000:10000),x(5000:10000,3),’LineWidth’,2); % SIM

plot(t(5000:10000),x(5000:10000,4),’LineWidth’,2); % CCS52

plot(t(5000:10000),x(5000:10000,7),’LineWidth’,2); % KRP

plot(t(5000:10000),x(5000:10000,11),’LineWidth’,2); % SK

xlabel(’Time (a.u.)’);

ylim([0 inf])

xlim([500 1000])

set(gca,’xticklabel’,[])

legend(’CYCD-CDKB Free’,’SIM’,’CCS52’,’KRP’,’CYCD-CDKA Free’)

title(sprintf(’%s’,current))

set(gca,’fontsize’, 14,’FontName’,’Helvetica’);

set(gcf,’color’,’w’)

snapnow % to publish

return

function [dx_dt]

=larkinpnas(t,x,k11,k1,lp,lm,kssim,k31,k311,j3,k41,k4,j4,k7,j7,k8,j8,k9,j9,k10,j10,ko,ko2 11,ko21,jtfb,k22,k151,j15,k161,k1611,j16,k131,k1311,k141,k14,lcm,lcp,k21,k211,k2111,k12,k121,k1211,kdsim1,kdsim,cdhs,ccs52ko,cycd3ko)

%CycBt

dx_dt(1)=k11+k1.*x(8)-(k21 + k211.*x(4) +k2111 .* x(5)).*x(1);

%MPF

dx_dt(2)=k11+k1.*x(8)-lp.*x(2).*(x(3)-(x(1) - x(2)))+lm.*(x(1)-x(2))-(k21 + k211.*x(4) +k2111 .* x(5)).*x(1)+(kdsim1 + kdsim .* x(2))*(x(1)-x(2));

%SIM

dx_dt(3)=kssim-(kdsim1 + kdsim .* x(2)).*x(3);

%Cdh1

dx_dt(4)=((k31+k311.*x(5)).*(cdhs-x(4))./(j3+cdhs-x(4))-(k41.*x(11)+k4.*x(2)).*x(4)./(j4+x(4)))*ccs52ko;

%Cdc20a

dx_dt(5)=k7.*x(6).*(1-x(5))./ (j7+ 1 - x(5)) - k8 .* x(5) ./ (j8 + x(5));

%IE

dx_dt(6)=k9 .*(1-x(6)).*x(2) ./ (j9 + 1-x(6)) - k10 .* x(6) ./ (j10 + x(6));

%CKIT

dx_dt(7)=ko - (k12 + k121.*x(11) + k1211 .* x(2)) .* x(7);

%TFB

dx_dt(8)=(ko211 + ko21 .* x(2)) .* (1-x(8)) ./ (jtfb + 1-x(8)) - k22 .* x(8) ./ (jtfb + x(8));

%TF

dx_dt(9)=k151 .* (1-x(9)) ./ (j15 + 1 - x(9)) - (k161 + k1611 .* x(11)) .* x(9) ./ (j16 + x(9));

%SKT

dx_dt(10)=(k131 + k1311 .* x(9))*cycd3ko - (k141 + k14.* x(2)).*x(10);

%SK

dx_dt(11)=(k131 + k1311 .* x(9))*cycd3ko - (k141 + k14.* x(2)).*x(11) + lcm .* (x(10) - x(11)) - lcp .* x(11) .* (x(7) - (x(10) - x(11))) +(k12 + k121.*x(11) + k1211 .* x(2))*(x(10)- x(11));

dx_dt=dx_dt’;

return

